# Skull allometries in three species of *Didelphis* (Didelphimorphia, Didelphidae)

**DOI:** 10.1101/2022.06.14.496063

**Authors:** Pere M. Parés-Casanova

**Affiliations:** Catalonia, Spain

**Keywords:** *Didelphis albiventris*, *Didelphis marsupialis*, *Didelphis pernigra*, isometry, Marsupialia

## Abstract

As biological shape is usually complex and evolves on different constraints, it can be assessed using integrative methods such as geometric morphometrics. Allometric changes were analysed in three species of *Didelphis* genus (*D. albiventris* n=20, *D. marsupialis* n=82, *D. pernigra* n=35) by means of geometric morphometric techniques. A significant correlation between shape and size was found, suggesting an allometric change pattern for all three species studied. However, allometries appeared to be different between *D. marsupialis* and *D. pernigra*, both of which belong to different groups (the so-called black-eared group and the white-eared group, respectively). Results are consistent with taxonomic recognition at the group level and can help to elucidate phylogenetic relationships between these three *Didelphis* species.

## INTRODUCTION

Allometry is defined as the relation between the size of an organism and its shape (Adams and Nistri 2010). Developed by Huxley can be described by simple functional form y/x=c, where c is a constant and x and y are size measures in units of the same dimension. In contrast to traditional morphometric approach which uses linear distances and angles, geometric morphometric (GM) methods are based on Cartesian coordinates of measurement points (Zelditch, Swiderski, and Sheets 2004) (Adams and Nistri 2010). In GM, shape is the geometric feature of an object without any nuisance parameters (position, rotation and scale), while size is interpreted as Centroid Size (CS), which is defined as “the square root of the sum of squared distances of a set of landmarks from their centroid” (Zelditch et al. 2004). Thus, in GM, allometry can be expressed by multivariate regression of the shape coordinates on CS (or its logarithm) (Zelditch et al. 2004).

Marsupials undergo great changes during their postnatal ontogeny (Abdala, Flores, and Giannini 2001). The New World family Didelphidae includes living marsupial species whose youngsters are born after a short gestation period, when basic processes of craniofacial morphogenesis are still active, but, as litter must be immediately capable of breathing, suckling and holding on to the mother’s teats, important trophic structures such as the premaxillae, maxillae, palatine and dentary bones are already in the process of ossification (Abdala et al. 2001) (Astúa and Leiner 2008).

Although species of the genus *Didelphis* have been morphometrically analysed separately in several studies, few allometric studies have been performed to date, and most of them have been undertaken within a single species (Thomason, Russell, and Morgeli 1990) (Abdala et al. 2001) (Cáceres 2002) (Schimming et al. 2016) (Mohamed 2018). So, comparative interspecific studies of their allometric variation could allow a better assessment of their taxonomic status.

The Andean White-eared Opossum (*D. pernigra* Allen 1900) (Gardner (e*D*.) 2007) and the White-eared Opossum sensu stricto (*D. albiventris* Lund 1840) are included within the “white-eared opossum group”, together with *D*. imperfecta. The Black-eared Opossum, Common Opossum or Southern Opossum (*D. marsupialis* Linnaeus 1758) belongs to the “black-eared opossum group”, together with *D*. aurita (Gardner (e*D*.) 2007).

In this research, we used GM techniques to address the following subjects aimed at addressing taxonomic issues in *Didelphis marsupialis, Didelphis albiventris* and *Didelphis pernigra*:

1. Analysis of the relationship between size and shape.
2. Visualization of these allometric patterns, and
3. Comparison of these patterns.

Our findings represent a contribution to the study of growth models in marsupials and can serve as a baseline for comparisons with other populations and species of *Didelphis*, as well as for future ecomorphological investigations aimed at understanding developmental trajectories in this group.

## MATERIALS AND METHODS

### Samples

We examined a sample of 137 specimens of *Didelphis* spp. (*D. albiventris* n=20, *D. marsupialis* n=82, *D. pernigra* n=35) archived in the collections of the Departamento de Biología of the Universidad del Valle in Cali (Colombia) and Instituto de Ciencias Naturales of the Universidad Nacional de Colombia in Bogotá. Every specimen had been taxonomically identified to the species level and was collected in Colombia for other purposes. A complete list of specimens may be obtained upon request to the author. The two-colour phases (grey and black) of the white-eared group were not considered.

### Taking photographs and digitizing

Digital images of skulls were taken with a Nikon D1500 digital camera equipped with an 18-105 mm Nikon DX telephoto lens. The photographic record was carried out using a standardized and homologous skull position for all specimens (facing up ventrally). Each specimen was placed in the centre of the optical field, with ventral aspect oriented parallel to the image plane. High quality pictures (1.5 to 6 MB) were saved in jpg format and transferred to the computer. Then a set of 14 landmarks on the ventral aspect of skull were digitized using TpsDig software v.2.16 (Rohlf 2015) to obtain the x-y coordinates of all points. The landmarks chosen were present on all specimens and were considered to sufficiently represent the morphology of the ventral aspect of skull, e.g., landmarks 1 and 7-10 describe buccal shape, landmarks 2-3 and 12-14 delineate braincase shape, while landmarks 4-6 and 11 represent shape of the zygomatic arches (Figure 1). The selection of these landmarks was based on used choices in other morphometric studies in mammal skulls (Cardini and O’Higgins 2005) (Flores, Giannini, and Abdala 2006) (Jamniczky and Hallgrímsson 2009) (López-Aguirre, Pérez-Torres, and Wilson 2015). All images included a ruler for scale (10 mm). Landmarks were digitized only on the left side to avoid redundant information. Each specimen was then superimposed onto the shapes by a translation, rotation and scalation, establishing the measure of shape by means of Procrustes coordinates.

**Figure 1.**
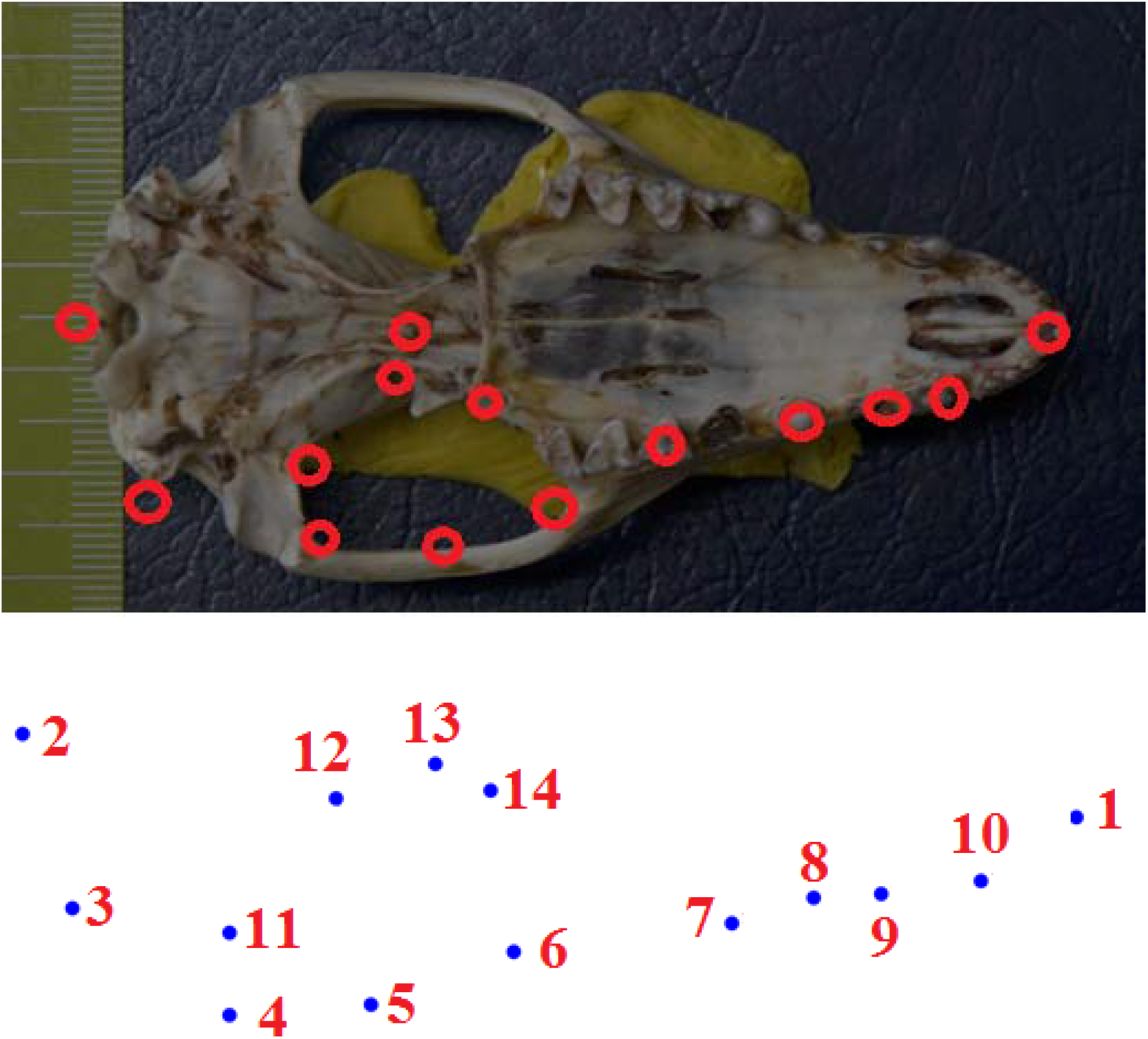
Location of the 14 landmarks used in the analysis. Fourteen landmarks were initially used to capture skull shape on the central skull view of *Didelphis* spp. (only on ventral left side). Skulls were aligned by their dorsal aspect to a stable plane. Landmarks 1 and 7-10 describe buccal shape, landmarks 2-3 and 12-14 delineate braincase shape, while landmarks 4-6 and 11 represent shape of the zygomatic arches. Landmarks 1, 2 and 13 are on the median line.

### Correlation between shape and tangent space

Correlations between the Cartesian points and tangent shape distances were calculated using TpsSmall software v. 1.29 (Rohlf 2015).

### Interaction with sex

With a NPMANOVA with species, sex, and sex by species interaction as independent variables analysed differences between species.

### Relationship between size and shape

Growth trajectories were analysed comparing trait changes against size, because animals’ information on age was not available. Centroid Size (CS) was interpreted as the measure of overall geometric size (Klingenberg 2016). Results are reported as a percentage value of the explained total shape variation from the predicted shape variation. These analyses were computed as permutation test with 10,000 runs and accompanied by shape deformation images to better display shape variations. To further quantify both intra- and interspecific differences on shape, we performed Canonical Variate Analysis (CVA), that is, a multivariate statistical method utilized to distinguish among multiple groups of specimens. The results of CVA are statistically reported as Mahalanobis distance (Md). Differences among the distances were determined by bootstrapping (10,000 runs). All analysis were performed with MorphoJ v. 1.06c software (Klingenberg 2011).

### Comparison of allometric patterns

We used the ordination analysis of the principal components (that is, Principal Component Analysis, PCA) to obtain the amount of variation, and the shape variation associated with first Principal Components (PC).

PCA was done from the var-covar matrix generated by the Procrustes coordinates, which includes the measures of the association between Procrustes coordinates themselves. We compared the allometry trajectories using the angular comparison between the corresponding vectors. This comparison has been done with the formula implemented in MorphoJ. Results are reported as angular values of the pairwise angular comparisons, with p-values for all the pairwise tests performed by the analysis.

## RESULTS

Variation of the specimens in shape space was perfectly correlated with tangent space for all anatomical aspects (r=0.997). This allowed us to use the plane approximation (skull ventral aspect) in subsequent statistical analyses. NPMANOVA reflected no differences among sexes (Table 1), so samples remain clustered for each species.

**Table 1.** NPMANOVA with species, sex, and sex by species interaction as independent variables to analyse species differences (*Didelphis marsupialis* n=72; *D. albiventris* n=18; *D. pernigra* n=34).

| Source | Sum of squares | Degrees of freedom | Mean square | F | *p* |
| --- | --- | --- | --- | --- | --- |
| Species | 5.39E+24 | 2 | 2.70E+24 | 2.8668 | 0.0087 |
| Sex | 6.28E+23 | 1 | 6.28E+23 | 0.6676 | 0.3676 |
| Interaction | 4.46E+24 | 2 | 2.23E+24 | 2.3732 | 0.0109 |
| Residual | 1.11E+26 | 118 | 9.40E+23 |  |  |
| Total | 1.21E+26 | 123 |  |  |  |

To assess both intra- and interspecific differences in the shape, CVA was performed between groups (i.e., population and species). If, on the one hand, this analysis highlighted the lack of significant interspecific variation only between the the “white-eared opossum group” (*D. albiventris* and *D. pernigra*) and the “black-eared opossum group” (*D. marsupialis*): *D. albiventris* vs. *D. pernigra*, Md=2.026; *D. albiventris* vs. *D. marsupialis*, Md=2.970; *D. pernigra* vs. *D. marsupialis*, Md=3.060.

Although the regression of Procrustes coordinates on log-CS accounted for a low variation for the three species (*D. pernigra*=16.99%; *D. albiventris*=27.93%; *D. marsupialis*=9.65%), the permutation test indicated the occurrence of significant ontogenetic allometries for each species (p<0.001) So the null hypothesis of isometry (a species that exhibits a constant shape regardless of size) was rejected (allometry was present) in all three species, since each multivariate regression was statistically significant. The regression trajectories are described for the three species by relating the CS log-transformed values vs. the shape scores (that is, the regression score obtained from the Procrustes coordinates) (Figures 2 to 4).

**Figure 2.**
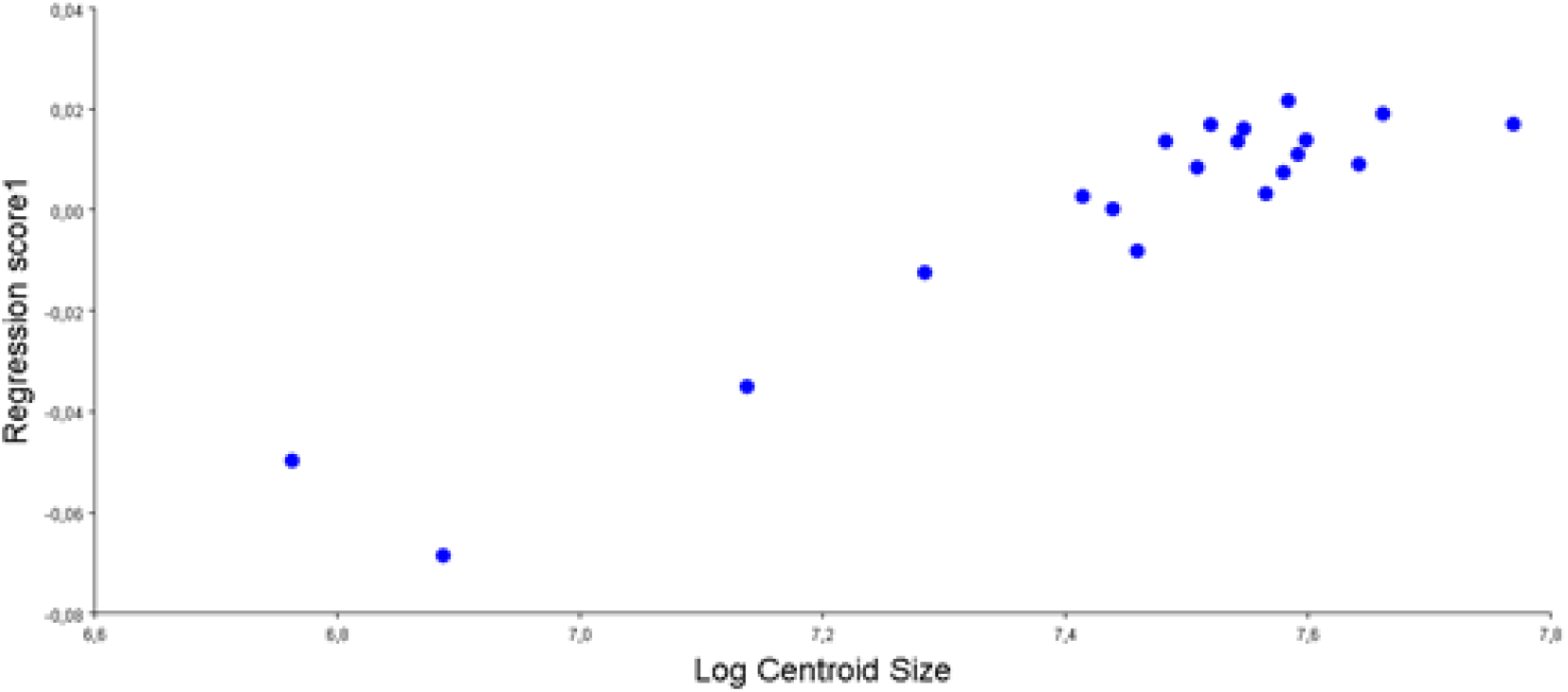
Regression of shape on centroid size (CS, log-transformed) for *Didelphis albiventris* (n=20). Figures represent the changes from smallest to biggest specimen. There was a significative change of shape according to size (p<0.001).

**Figure 3.**
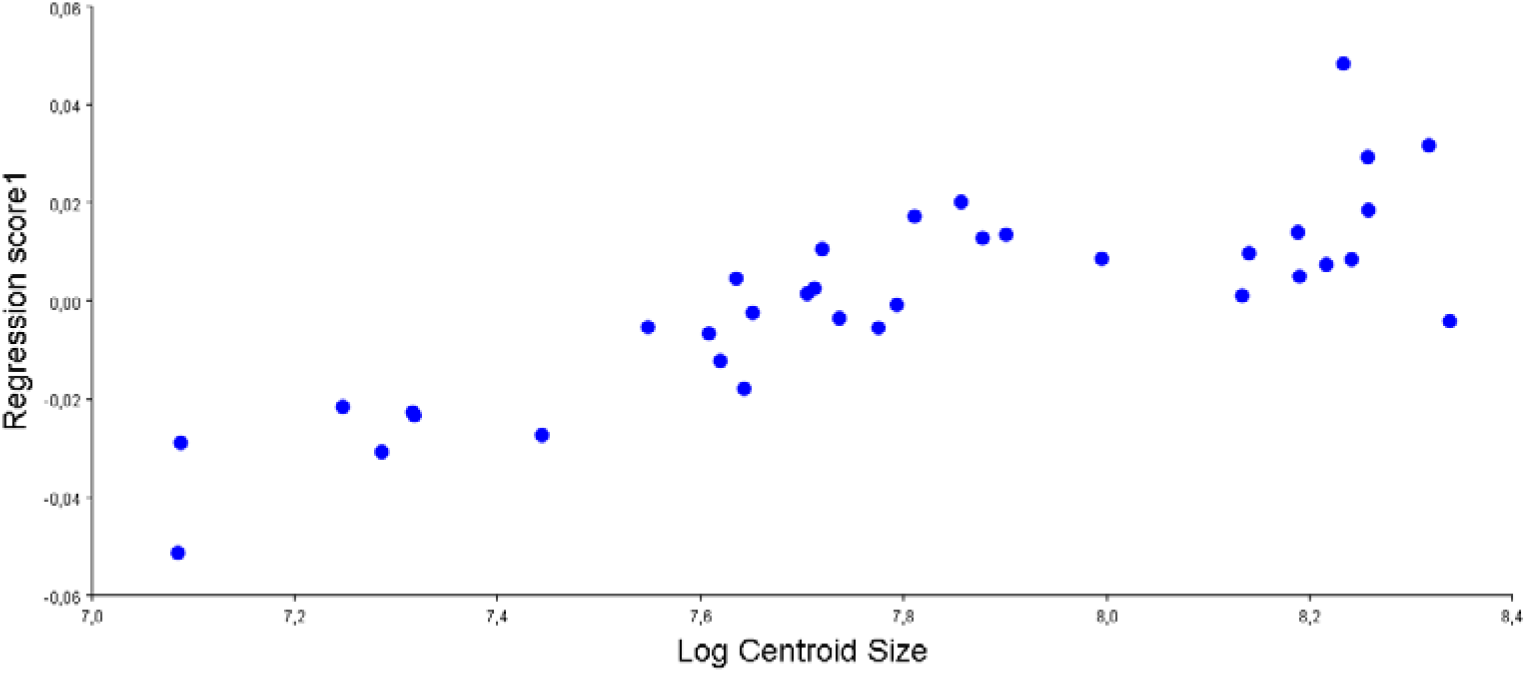
Regression of shape on centroid size (CS, log-transformed) for *D. pernigra* (n=35). Figures represent the changes from smallest to biggest specimen. There was a significative change of shape according to size (p<0.001).

**Figure 4.**
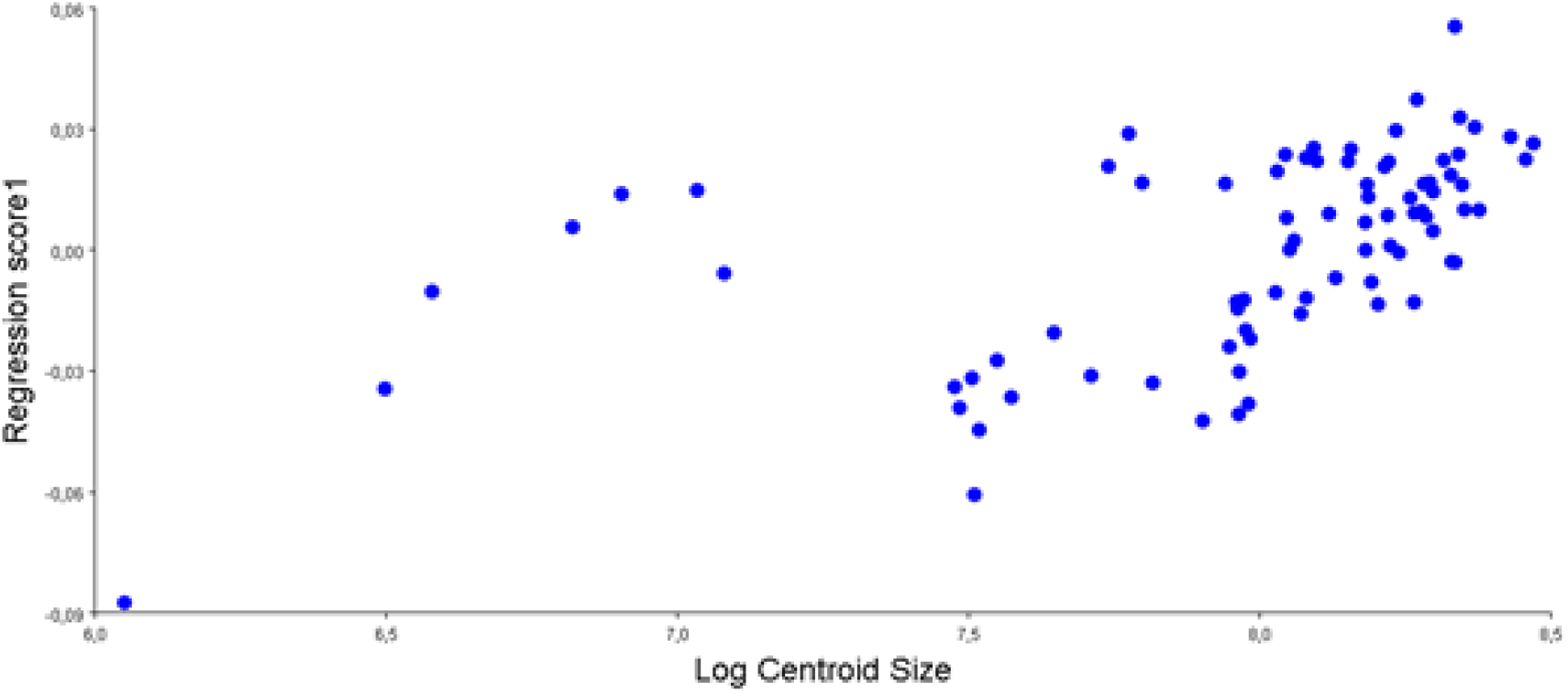
Regression of shape on centroid size (CS, log-transformed) for *Didelphis marsupialis* (n=82). Figures represent the changes from smallest to biggest specimen. There was a significative change of shape according to size (p<0.001).

The PCA scatter plot showed a high morphological variation of the entire sample. The first principal component (PC1) explained 28.78% of the total variance and PC2 explained 13.87% (Figure 5), while each of the 22 remaining PCs explained 57.35%, which is an important amount of shape variation information. Along the PC1 axis individuals varied mainly in relative length of neurocranium, while PC2 captured mainly the relative degree of basilar part of occipital bone, with the splanchnocranium being the most conservative skull region. Deformation grids of the axis’ extremes are showed in Figure 6.

**Figure 5.**
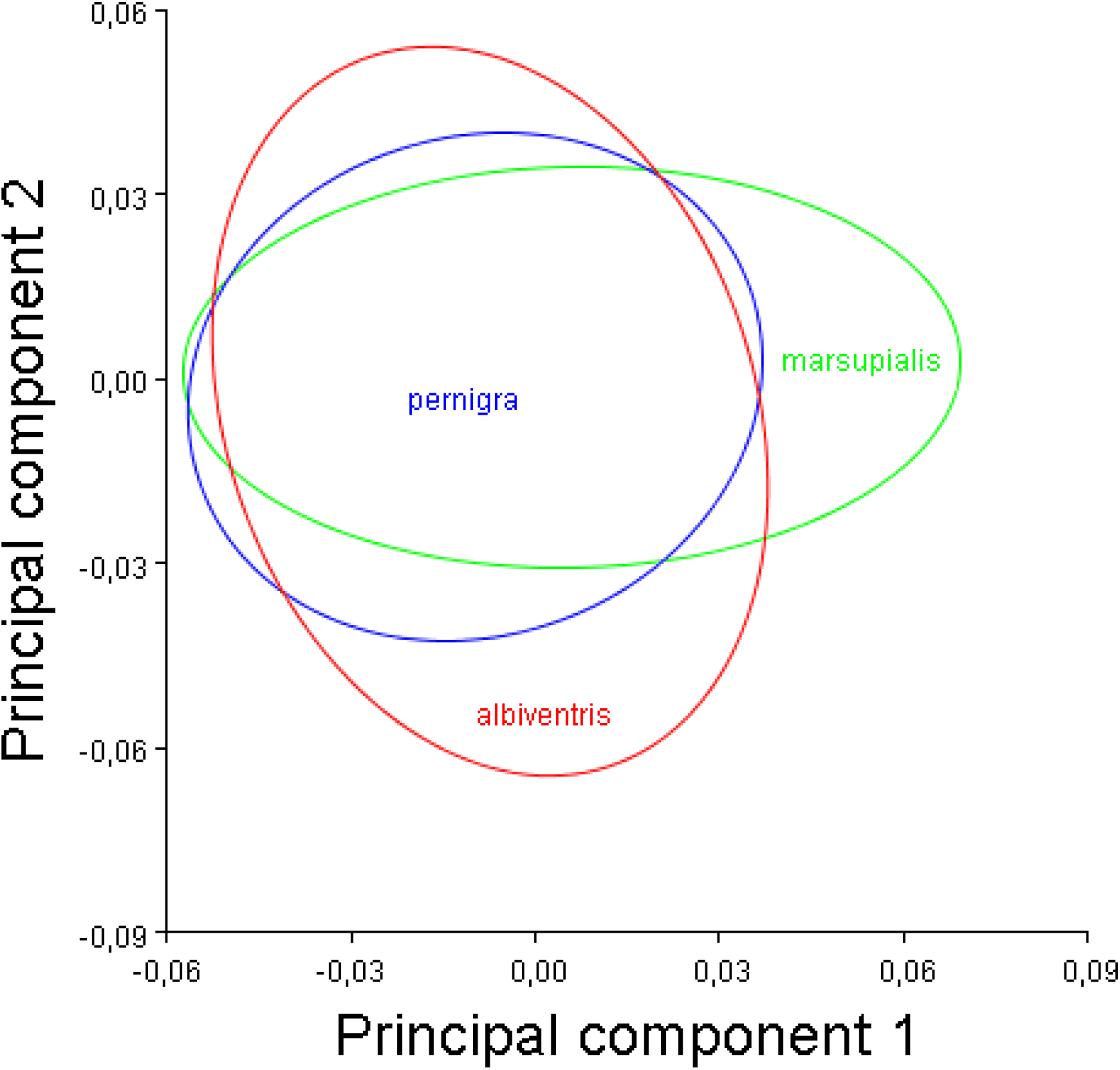
Principal Component Analysis of multivariate allometric patterns in *D. albiventris* (n=20), *D. marsupialis* (n=82) and *D. pernigra* (n=35). PC scores of the allometric patterns of each species and 90% confidence ellipses from the respective boostraps estimates are plotte*D*. The first PC explained 28.78% of the total variance and the second PC a 13.87%.

**Figure 6.**
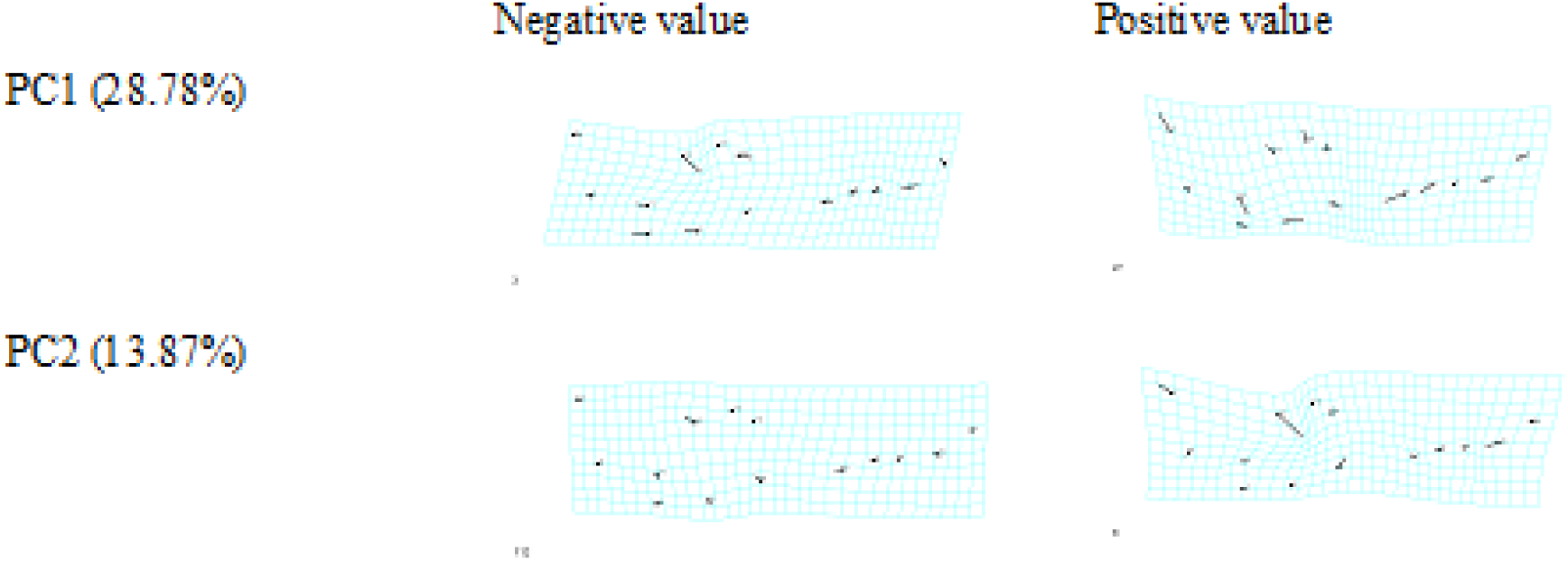
The skull shape variation of the whole sample. Deformation grids are associated to the most negative (left) and positive (right) values of the first two principal components (PC1 and PC2).

The angular comparisons between the three species showed non-significant distinctions between the ontogenetic trajectories of *D. albiventris* vs. *D. pernigra*=86.34 (p=0.762); and *D. albiventris* vs. *D. marsupialis*=73.79 (p=0.176), but for *D. pernigra* vs. *D. marsupialis*=29.61 (p<0.0001).

Shape changes associated with size increase showed similar variation patterns between *D. albiventris* and *D. marsupialis*, but there appeared some differences between the latter two and *D. pernigra*. Shape variations can be described as follow: in all the species, body shapes were focused on braincase, although this was more centred on basilar region in *D. pernigra*. Then, the main shape changes involved the braincase, the face seeming to a more conservative region.

## DISCUSSION

Although most of developmental shape changes are known to occur earlier in ontogeny, *Didelphis* is known to continue growing after full eruption of dentition (Astúa 2015) and this fact would explain why allometry accounted for a low but significant amount of postnatal ontogenetic shape variation. This differentiation would be centered on the central nervous system and sensory capsules, thus producing more change in braincase, eyes, and auditory region (neurocranial components) in comparison to trophic components (splanchnocranial components) of the skull. As in marsupials, neurogenesis occurs after birth and during lactation (Abdala et al. 2001), this conclusion is not surprising.

Finally, results presented here are therefore consistent with taxonomic recognition at the group level (“white-eared” and “black-eared”), as no different trajectories appeared in the two closely related species of the white-eared group (*D. albiventris* and *D. pernigra*). The allometric difference with *D. marsupialis*, moreover, would corroborate *D. pernigra* as a different species of *D. albiventris*, thus reinforcing the logics of the “white eared group”.

## ACKNOWLEDGEMENTS

I thank Catalina Cárdenas and Hugo Fernando López, from *Instituto de Ciencias Naturales* of the *Universidad Nacional de Colombia* in Bogotá (Colombia), and Óscar Murillo, from *Departamento de Biología* of the *Universidad del Valle* in Cali (Colombia), for access to their institutional collections. No financial assistance was received.

## REFERENCES

Abdala, F., D. A. Flores, and N. P. Giannini. 2001. “Postweaning Ontogeny of the Skull of *Didelphis albiventris*.” Journal of Mammalogy 82(1):190–200.

Adams, D. C. and A. Nistri. 2010. “Ontogenetic Convergence and Evolution of Foot Morphology in European Cave Salamanders (Family: Plethodontidae).” BMC Evolutionary Biology 10:216.

Astúa, D. 2015. “Morphometrics of the Largest New World Marsupials, Opossums of the Genus Didelphia (Didelphimorphia, Didelphidae).” Oecologia Australis 19(1):117–42.

Astúa, D. and Natália Oliveira N. O. Leiner. 2008. “Tooth Eruption Sequence and Replacement Pattern in Woolly Opossums, Genus Caluromys (Didelphimorphia: Didelphidae).” Journal of Mammalogy 89(1):244–51.

Cáceres, Nilton C. 2002. “Food Habits and Seed Dispersal by the White-Eared Opossum, *Didelphis albiventris*, in Southern Brazil.” Studies on Neotropical Fauna and Environment 37(2):97–104.

Cardini, A. and P. O’Higgins. 2005. “Post-Natal Ontogeny of the Mandible and Ventral Cranium in Marmota Species (Rodentia, Sciuridae): Allometry and Phylogeny.” Zoomorphology 124(4):189–203.

Flores, D. A., N. P. Giannini, and F. Abdala. 2006. “Comparative Postnatal Ontogeny of the Skull in the Australidelphian Metatherian *Dasyurus Albopunctatus* (Marsupialia: Dasyuromorpha: Dasyuridae).” Journal of Morphology 267(February):426–40.

Gardner (eD.), A. L. 2007. “Marsupials, Xenarthrans, Shrews, and Bats.” P. 690 in Mammals of South America, vol. 1, edited by The University of Chicago Press. Chicago.

Jamniczky, Heather A. and B. Hallgrímsson. 2009. “A Comparison of Covariance Structure in Wild and Laboratory Muroid Crania.” Evolution 63(6):1540–56.

Klingenberg, C. P. 2011. “MorphoJ: An Integrated Software Package for Geometric Morphometrics.” Molecular Ecology Resources 11(2):353–57.

Klingenberg, C. P. 2016. “Size, Shape, and Form: Concepts of Allometry in Geometric Morphometrics.” Development Genes and Evolution 226(3):113–37.

López-Aguirre, C., J. Pérez-Torres, and L. A. B. Wilson. 2015. “Cranial and Mandibular Shape Variation in the Genus *Carollia* (Mammalia: Chiroptera) from Colombia: Biogeographic Patterns and Morphological Modularity.” PeerJ 3(8):1–23.

Mohamed, R. 2018. “Anatomical and Radiographic Study on the Skull and Mandible of the Common Opossum *(Didelphis marsupialis* Linnaeus, 1758) in the Caribbean.” Veterinary Sciences 5(44):1–10.

Rohlf, F. J. 2015. “The Tps Series of Software.” Hystrix 26(1):9–12.

Schimming, B. C., L. F. F. Reiter, L. M. Sandoval, L. André, L. R. Inamassu, and M. J. Mamprim. 2016. “Anatomical and Radiographic Study of the White-Eared Opossum (*Didelphis albiventris*) Skull.” Pesq. Vet. Bras. 36(11):1132–38.

Thomason, J. J., A. P. Russell, and M. Morgeli. 1990. “Forces of Biting, Body Size, and Masticatory Muscle Tension in the Opossum *Didelphis virginiana*.” Canadian Journal of Zoology 68(2):318–24.

Zelditch, M. L., D. L. Swiderski, and H. D. Sheets. 2004. Geometric Morphometrics for Biologists: A Primer. Boston: Elsevier Academic Press.

